# An AAV-ie based Vaccine effectively protects against SARS-CoV-2 and Circulating Variants

**DOI:** 10.1101/2021.05.19.444889

**Authors:** Simeng Zhao, Fangzhi Tan, Junzi Ke, Jie Yang, Chao-bo Lin, Haopeng Wang, Guisheng Zhong

## Abstract

Prophylactic vaccines against SARS-CoV-2 have been extensively developed globally to overcome the COVID-19 pandemic. However, recently emerging SARS-CoV-2 variants B.1.1.7 and B.1.351 limit the vaccine protection effects and successfully escape antibody cocktail treatment. Herein, based on our previously engineered adeno-associated viral (AAV) vector, AAV-ie, and systematic immunogen screening, we developed an AAV-ie-S1 vaccine with thermostability, high efficiency, safety, and single-dose vaccination advantage. Importantly, the AAV-ie-S1 immune sera efficiently neutralize B.1.1.7 and B.1.351, indicating a potential to circumvent the spreading of SARS-CoV-2.

---

Severe acute respiratory syndrome coronavirus-2 (SARS-CoV-2) has caused the COVID-19 pandemic, with more than 117 million infections and 2.5 million deaths. To fight against this rapidly spreading pandemic, prophylactic vaccines have been developed using different techniques, such as inactivated virus,^1–3^ messenger RNAs,^4,5^ recombinant proteins,^6^ and viral-vectored vaccines.^7,8^

However, rapidly spreading variants of SARS-CoV-2 have been emerging. Mutations in spike (S) protein have raised deep concerns of vaccine efficacy since the S protein is the main target of vaccines and neutralizing antibodies. Recent studies have shown that the efficiency of mRNA vaccines and inactivated vaccines were significantly decreased against newly occurred variants with E484K mutation, such as B.1.351 and B.1.1.248,^9–13^ suggesting the urgent need for effective vaccines against SARS-CoV2 circulating variants.

Adeno-associated viruses (AAVs) belong to a class of non-enveloped single-stranded DNA dependoparvovirus, and have been widely used as gene delivery vectors because of their efficacy and low immunogenicity^14^. To date, three AAV-based gene therapies have been approved, including Luxturna (inherited retinal diseases, IRD), Zolgensma (spinal muscular atrophy, SMA), and Glybera (familial lipoprotein lipase deficiency, LPLD). AAVs were also proven to be promising vaccine vectors.^15,16^ We initiated the development of AAV-vectored COVID19 vaccine candidates based on our engineered AAV variant, AAV-ie,^17^ and explored the potential of this AAV as a vaccine platform.

Our earlier study showed that AAV-ie effectively infects many types of cochlear cells, likely due to the insertion of the membrane-permeable peptide.^17^ We hypothesized that AAV-ie might have a broad tropism, including muscles, the main targeted infection sites for vaccines.^18^ We first prepared an AAV-ie vector expressing nuclear located GFP reporter (AAV-ie-GFP) and investigated the tissue tropism of AAV-ie by intramuscular (i.m.) or intravenous (i.v.) injecting AAV-ie-GFP at the dose of 1 × 10^11^ genome copies (GCs) per mouse (1 × 10^12^ GC/mL, 100 μL). Tissue imaging results showed that intramuscular (i.m.) AAV-ie injection resulted in highly transduced muscle cells around the injecting site. In contrast, almost no GFP expression was detected in other organs, including the brain (Extended Data Fig. 1a), suggesting that i.m. AAV-ie injection resulted in a highly effective and specific expression of targeted proteins in **muscles**. Intravenous (i.v.) resulted in high level of expression of GFP in livers and hearts. Low extent infection was also observed in spleens (Extended Data Fig. 1b). Besides, neither i.v. nor i.m. injection of AAV-ie resulted in body weight loss (Extended Data Fig. 1c). Our results support that AAV-ie may act as an appropriate vector for SARS-CoV-2 vaccine development.

We then generated a series of vaccine candidates expressing different SARS-CoV-2 S domains with an IL-2 leading signal (Extended Data Fig. 2a). To mimic the trimerized natural S protein, in some cases, a C-terminal T4 fibritin domain (Fd) was fused to stabilize trimer formation. All candidates induced antigen expression in HEK293T cells (Extended Data Fig. 2b). We then vaccinated the mice with the candidates at the dose of 1 × 10^11^ GCs per mouse by i.v. injection and monitored the S-binding antibody titers. Injection of AAV-ie-S1 induced the highest IgG level (Extended Data Fig. 2c-2d) and a balanced IgG2a/IgG1 ratio (Extended Data Fig. 3). Thus, S1 was chosen as the immunogen for further investigations.

We vaccinated the mice with AAV-ie-S1 by i.m injection at the dose of 6 × 10^10^ GCs per mouse and monitored the antibody titers. AAV-ie-S1 stimulated robust humoral responses, with the S-binding geometric mean titers (GMTs) as 3.18 × 10^4^, 3.54 × 10^5^, 2.06 × 10^6^, 2.55 × 10^4^, and 3.83 × 10^4^ on week 4, 6, 8, 12, and 16, respectively (Extended Data Fig. 4a-4b). The RBD-binding antibody level was evaluated and the GMTs were 1.33 × 10^4^, 1.15 × 10^5^, 9.58 × 10^4^, 1.10 × 10^5^, and 5.34 × 10^4^ on week 4, 6, 8, 12, and 16, respectively (Extended Data Fig. 4c-4d). The immune sera were tested for their pseudo-virus neutralizing activity, and the neutralizing GMT reached the peak value at week 8 (EC_50_ = 137), consistent with the binding GMT (Extended Data Fig. 5). Moreover, AAV-ie-S1 was tested for its thermostability by vaccinating the mice with the vaccine stored at different temperatures. Following antibody titer detection showed that AAV-ie-S1 stimulated a similar immune response after stored for 2 weeks at 4 °C or even room temperature (Extended Data Fig. 6).

To evaluate the potential of AAV-ie-S1 as a vaccine against SARS-CoV-2, we tested its immunogenicity in the *Macaca fascicularis*. Four macaques were divided into two groups and treated with AAV-ie-S1 or control AAV (AAV-ie-GFP) by i.m. injection at the dose of 1 × 10^13^ GCs per monkey. High S-binding antibody levels were observed in both macaques on week 4 after vaccination and were steadily maintained in the following two months (Fig. 1a-1c). The RBD-binding antibody titers were tested, and similar results were observed (Extended Data Fig. 7). The pseudo-virus neutralizing EC_50_ values of monkey 1 immune sera were determined as 236, 482, and 337 on week 4, 8, and 12, respectively; the neutralizing EC_50_ values of monkey 2 immune sera were determined as 1017, 1040, and 638 on week 4, 8, and 12, respectively (Fig. 1d-1f). These results showed that AAV-ie-S1 could induce robust humoral responses in non-human primates (NHPs).

**Fig. 1.**
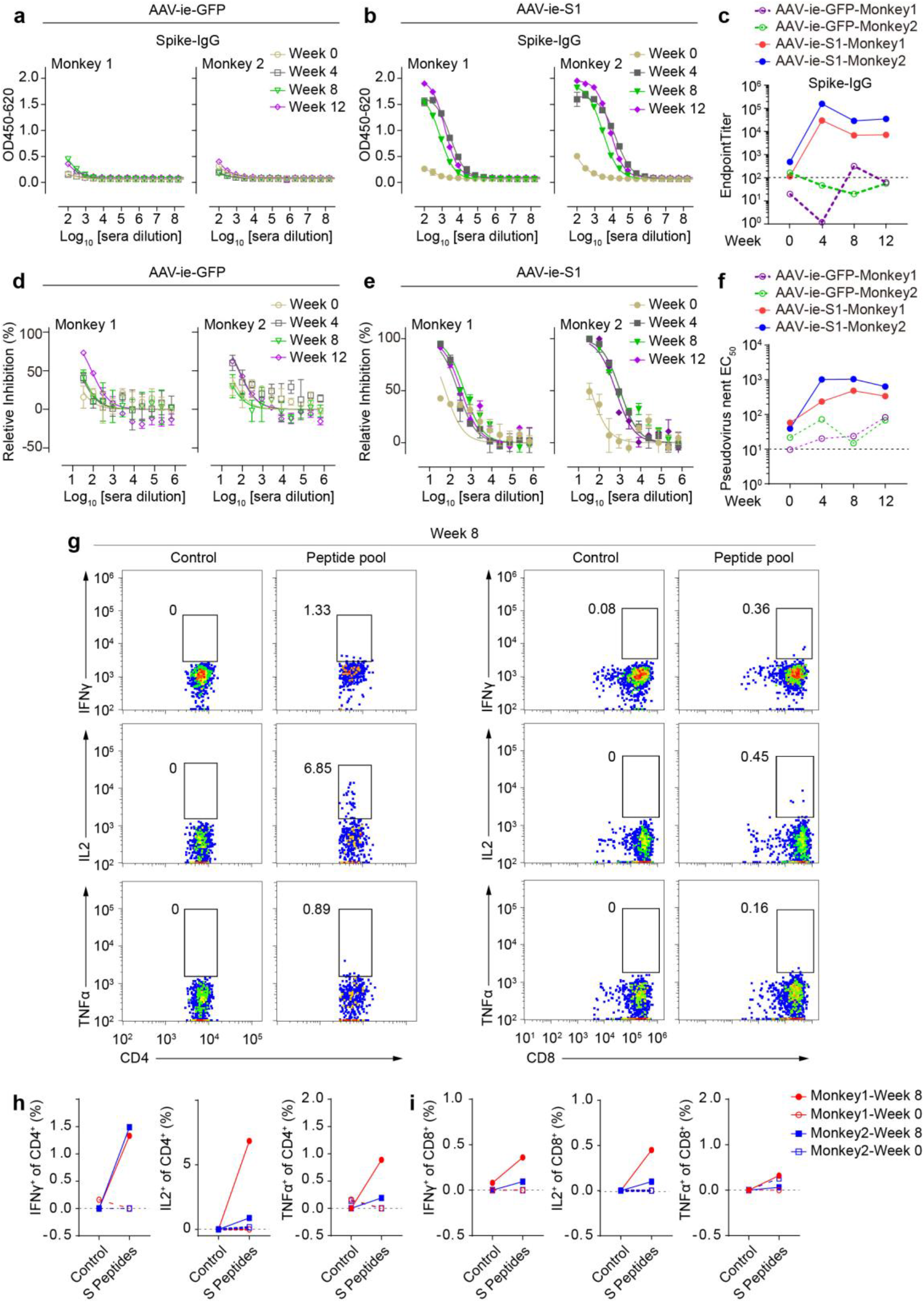
AAV-ie-S1 induces humoral and cellular responses against SARS-CoV-2 in NHPs. **a-c,** S-Binding IgG antibody titers of monkey sera after vaccination with AAV-ie-GFP (**a**) and AAV-ie-S1 (**b**) at indicated time. Endpoint titers were summarized in **c**. Experiments were performed in triplicates. **d-f**, SARS-CoV-2 pseudo-virus neutralizing activities of monkey immune sera vaccinated with AAV-ie-GFP (**d**) and AAV-ie-S1 (**e**) at indicated time. The neutralizing EC_50_ values were summarized in **f**. Experiments were performed in triplicates. **g**, Flow cytometry panels of cytokine staining of PBMCs after vaccination (week 8). PBMCs were stimulated with S peptide pool or vehicle control, and following intracellular staining of IFNγ, IL2, and TNFα was performed. **h-i**, Populations of CD4^+^ (**h**) and CD8^+^ (**i**) T cells responding to SARS-CoV-2 S peptides before (week 0) and after (week 8) vaccination using intracellular staining of indicated cytokines.

We collected peripheral blood mononuclear cells (PBMCs) from monkeys before (week 0) and after (week 8) vaccination to evaluate **T cellular responses** by stimulating the PBMCs with an S peptide pool. Compared to week 0, the IFNγ^+^, IL2^+^, and TNFα^+^ populations of CD4^+^ T and CD8^+^ cells were upregulated in week 8 PBMCs (**Fig. 1g-1i**, Extended Data Fig. 8). These results indicated that AAV-ie-S1 induced S-specific T cell responses after vaccination.

Recently, SARS-CoV-2 variants B.1.1.7 and B.1.351 are rapidly spreading and resistant to cocktail neutralizing antibody treatments and different types of vaccines^9–13^. The extensive mutations in the spike of B.1.1.7 and B.1.351 may cause antigenic changes that limit antibody treatment efficiency and vaccine protection. We evaluated the neutralizing efficiency of AAV-ie-S1 against B.1.351 and B.1.1.7. The FACS based S-binding assay was used to detect the binding abilities of immune sera of NHPs (week 12) to S proteins of wild-type (WT) **and mutated SARS-CoV-2 strains** (Extended Data Fig. 9a). The S-binding EC_50_ values of monkey 1 immune serum to WT, B.1.351 and B.1.1.7 S proteins were determined as 1.26 × 10^4^, 1.18 × 10^4^, and 1.54 × 10^4^, respectively; and the EC_50_ values of monkey 2 immune serum were 4.56 × 10^4^, 4.15 × 10^4^, and 5.83 × 10^4^, respectively (Fig. 2a-2c). These results showed that AAV-ie-S1 immune sera effectively bind to S proteins of WT SARS-CoV-2, B.1.1.7 and B.1.351. We also tested the binding abilities of mice immune sera, similar to NHPs, the binding activities of mice sera were not significantly affected by the mutations (Extended Data Fig. 9b).

**Fig. 2.**
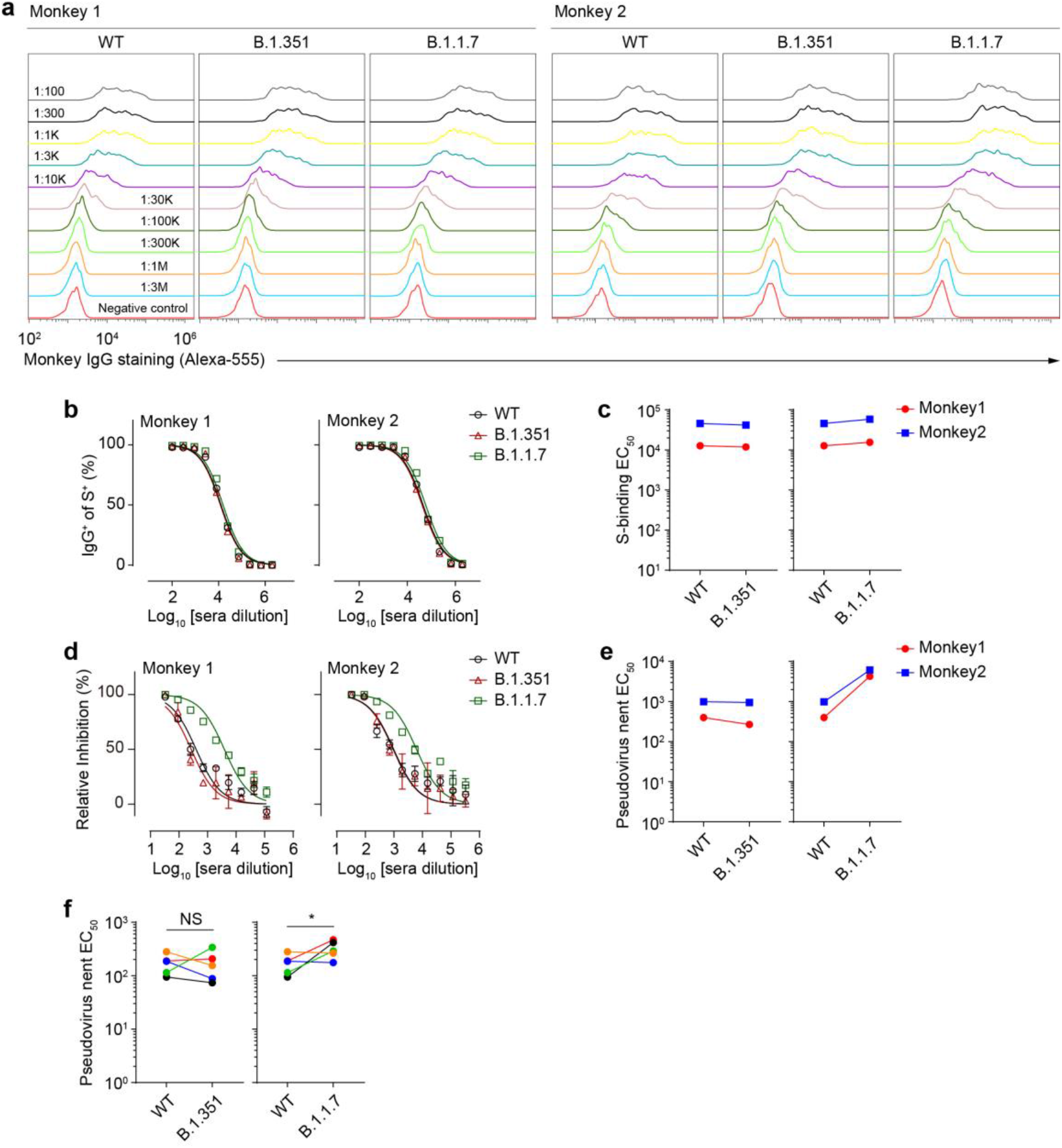
AAV-ie-S1 immune sera avoid mutational immune escape. **a**, Binding of serially diluted immune sera to S proteins of wild type, B.1.352, and B.1.1.7 SARS-CoV-2 variants expressed on the surface of HEK-293T cells. **b**, Summary of IgG^+^ populations under serially diluted immune sera staining. **c**, S-binding EC_50_ values summary of immune sera. **d-e**, Pseudo-virus neutralizing activities of monkey immune sera against WT SARS-CoV-2 and circulating variants. Dose curves were shown in (**d**) and the neutralizing EC_50_ values were summarized in **e**. Experiments were performed in triplicates. **f**, Neutralizing EC_50_ values of mice immune sera against WT SARS-CoV-2 and circulating variants. NS: not significant; * p < 0.05. N = 5 mice, experiments were performed in triplicates.

The pseudo-virus neutralizing experiments were then carried out to determine the potential efficacy of AAV-ie-S1 immune sera to neutralize SARS-CoV-2 variants B.1.351 and B.1.1.7. Intriguingly, compared to WT, the neutralizing abilities of NHPs immune sera against B.1.351, which has been reported to be resistant to first generation vaccines^9–13^, were not significantly decreased (EC_50_ values: 397 *v.s.* 267 for monkey 1; 985 *v.s.* 943 for monkey 2) (Fig. 2d-2e). Interestingly, the neutralizing activities against B.1.1.7 were 6 to 10 times higher (EC_50_ values: 397 *v.s.* 4261 for monkey 1; 985 *v.s.* 6104 for monkey 2) (Fig. 2d-2e). We also tested the neutralizing efficiency of mice immune sera to these circulating variants. Similar to the monkey immune sera, no significant immune escape was observed for B.1.351 variant. Mean neutralizing EC_50_ values against WT and B.1.351 were determined as 172 and 172, respectively (Fig. 2f). Meanwhile, the neutralizing activity against B.1.1.7 was 1.8 fold increased, and mean neutralizing EC_50_ was determined as 323 (Fig. 2f). These observations were consistent with the results of NHPs immune sera. Collectively, these results support that AAV-ie-S1 vaccine holds the potential to fight against the SARS-CoV-2 variants B.1.351 and B.1.1.7.

In summary, we developed a new type of SARS-CoV-2 vaccine based on an engineered AAV vector. A recent study also showed that AAVrh32.22 based vaccines encoding full-length S or S1 could stimulate robust humoral and cellular responses. However, whether this vaccine avoids mutational immune escapes remains unclear.^19^ Our results suggest that S1 is a better immunogen for SARS-CoV-2 vaccine development than the full-length S protein. AAV-ie-S1 vaccine showed several advantages, such as thermostability, high efficiency, safety, and single-dose vaccination. More importantly, AAV-ie-S1 vaccinated sera avoided mutational immune escape and neutralized emerging variants, such as B.1.351, efficiently.^9–13^. To advance our vaccine to the clinical stage, more experiments are needed. Our strategies of designing a vaccine may provide a novel avenue for future vaccine development to circumvent the spreading of SARS-CoV-2.

## Methods

### Vaccine design and production

Different domains of SARS-CoV-2 S protein were expressed under the control of the CMV promoter and terminated by an SV40 polyadenylation signal.

Research-grad and small-scale AAV vectors were homemade. Vector preparations were generated by calcium phosphate transfection of adenoviral helper plasmid, AAV-ie capsid construct, and ITR-flanked transgene plasmid in HEK293T cells. Cells and medium were harvested 96 hours after transfection. The collected cells then were treated with chloroform and the supernatant was collected. The supernatant and the medium were combined and concentrated with 1M NaCl and 10% PEG8000. After centrifugation, the pellet was resuspended in PBS buffer with DNase I (Thermo Fisher Scientific, Cat# EN0521) and RNase A (Tiangen, Cat# RT405). The crude AAVs were overlaid to iodixanol gradient solutions. After centrifugation, the AAV-containing 40% fraction was collected and concentrated. The genome-containing titers of vectors were determined by SYBR (Vazyme, Cat# Q311) analysis using primers targeting the WPRE region. The qPCR primers for WPRE are listed as follows: forward, 5’-CTTGTTTATTGCAGCTTATAATGG-3’; reverse, 5’-GATACATTGATGAGTTTGGACAAAC-3’.

### *In vitro* antigen expression and detection

Six-well plates seeded with HEK293T cells were transfected with different SARS-CoV-2 S truncated proteins expressing vectors (2μg DNA/plasmid) using the calcium phosphate transfection method. The supernatants were harvested 48 h post-transfection and mixed with reducing sample buffer (CWBio, Cat# CW0027S), heated for 8 minutes at 95°C and run on a precast 4-12% Bis-Tris PAGE gel (Genscript, Cat# M00654). Protein was transferred to a nitrocellulose (NC) filter membrane, and the membrane was blocked overnight at 4°C in PBS buffer containing 0.1% Tween 20 (PBST) and 5% non-fat milk powder. Then, the NC membrane was incubated for 1 hour in 5% milk PBST with a 1:1,000 dilution of mouse anti-His tag antibody (Proteintech, Cat# 66005-1-lg). After this, the NC membrane was washed three times with PBST buffer and subsequently incubated with 1:10,000 HRP-conjugated goat anti-mouse IgG (Proteintech, Cat# SA00001-1) in 5% milk PBST. Finally, the NC membrane was rewashed three times with PBST and imaged with ECL Western blotting detection system (Biorad, Cat# 1705062).

### Protein expression

Recombinant SARS-CoV-2 S and RBD proteins used for ELISA binding were self-expressed and purified. The coding sequences of S-trimer ECD (residues 1-1208) or RBD were cloned into pTT5 vector with a N-terminal IL-2 signal peptide and a C-terminal 6 × His tag. For S-trimer expression, four substitutions at the cleavage site (residues 682-685) were changed to “GSAS”, two proline substitutions at the residues 986 and 987 were also introduced to stabilize the protein as previously reported^20^. The plasmids were transfected into HEK Expi293F cells (Thermo Fisher Scientific, Cat# A14527) using polyethylenimine (PEI) method. Supernatants were collected after 5 days culture, and proteins were purified with Ni-NTA resin (Qiagen, Cat# 151010181) and followed by Superose 6 10/300 gel filtration column chromatography (GE Healthcare, Cat# 17-5172-01). Proteins were then concentrated and the concentrations were determined using BCA protein concentration kit (Pierce, Cat# 23225).

### Immunofluorescence

Mice were i.m. or i.v. injected with AAV-ie-GFP at the dose of 1 × 10^11^ GCs per mouse. After 2 weeks, mice were anesthetized with sodium pentobarbital (40 mg/kg, i.p.) with no avoidance response to foot pinch. They were then perfused with normal saline (at 37 °C) and subsequently by ice-cold 4% PFA for fixation. Organs were post-fixed in 4% PFA for 4 h, then dehydrated in 30% sucrose. organs were frozen at −80 °C and then sectioned at 20-μm-thick with the freezing microtome (Leica CM1950) for immunofluorescence labeling. Organs sections were rinsed in PBS, permeabilized, and then blocked with blocking solution (3% w/v donkey serum and 0.5% v/v Triton X-100 in PBS) for 1.5 h at room temperature. The slices were then stained using rabbit anti-GFP antibody (Proteintech, Cat# 50430-2-AP) in 0.1% Triton X-100 and 1% serum in PBS overnight at 4 °C. After washing with PBS, sections were incubated with the Alexa-488 conjugated donkey anti-rabbit antibody (Thermo Fisher Scientific, Cat# A21206) (1:1000 dilution) for 2 h at room temperature. After incubation, sections were washed with PBS. Sections were mounted with VECTASHIELD mounting medium for fluorescence with DAPI (Vectorlab, Cat# H1200). Following imaging was performed using confocal microscope (Nikon)

### Mouse vaccination

The mouse study was performed under the guidance of Institutional Animal Care and Use Committee (IACUC) of Shanghaitech University, China. Male BALB/c mice aging 6 – 8 weeks were used in this study. Mice were housed under a 12 h light/dark cycle and allowed free access to diet and water. For primary antigen screening, mice were i.v. injected with of the AAV vectored vaccines at the dose of 1 × 10^11^ GCs per mouse, after 2 weeks, the blood samples were collected from the retro-orbital plexus to prepare immune sera. For AAV-ie-S1 vaccination, the vaccine was i.m. injected at the dose of 6 × 10^10^ GCs per mouse. The blood was then collected in regular intervals to prepare immune sera. Sera were kept at −80 °C before use.

### Non-human primate (NHP) vaccination

All animal procedures were approved by the Institutional Animal Care and Use Committee of Shanghaitech University. *Macaca fascicularis* (2 to 5 year-old) that screened negative for viral pathogens were enrolled on the study. Animals were housed in in stainless-steel squeeze back cages, on a 12-hour timed light/dark cycle, at room temperatures. NHPs were treated with varied enrichments such as food, visual and auditory stimuli, and social interactions throughout the study. Four *Macaca fascicularis* were treated with the vaccine candidates (N = 2 males in each group) intramuscularly at a dose of 1 × 10^13^ GCs per animal (1 × 10^13^ GCs/mL, 1mL). Sera samples were obtained in regular intervals to analyze the immunogenicity of AAV-ie-S1. And whole blood samples were collected before (week 0) and after (week 8) vaccination to isolate PBMCs for cellular response analysis.

### ELISA

To evaluate the antibody titers binding to SARS-CoV-2 S or RBD protein in the sera of immunized animals. Nunc Maxisorp plates (Thermo Fisher Scientific, Cat# 464718) were coated with recombinant S or RBD protein at the concentration of 2 μg/mL in PBS at 4 °C overnight. After extensive washing with PBS, the plates were blocked using 5% skim milk at RT for 2 h. Serially diluted immune sera of vaccinated mice or monkeys were then added to the plates and incubated at RT for 1 h. After washes with PBS, HRP-conjugated secondary antibodies (HRP conjugated goat anti-mouse IgG: Proteintech, Cat# SA00001-1; HRP conjugated goat anti-mouse IgG2a: Proteintech, Cat# SA00012-2; HRP conjugated goat anti-mouse IgG1: Proteintech, Cat# SA00012-1; HRP-conjugated goat anti-monkey IgG: Southern Biotech, Cat# 4700-05) were added and incubated for 1 h. The plates were further washed using PBS and TMB substrate (Beyotime, Cat# P0209) was added. The absorbance at 450/620 nm was then measured with a micro plate reader (Flexstation III, Molecular devices).

### Pseudo-virus neutralization assay

To prepare SARS-CoV-2 and related mutated pseudo-viruses, HEK-293T cells were co-transfected with pcDNA3.1-SARS-CoV-2-S or related mutants and pNL4-3.luc.RE by calcium phosphate transfection. After 72 h culture, the supernatant was collected and centrifuged at 3,000 X *g* for 10 min. To test the neutralizing activity of immune sera against pseudo-virus, HEK293T cells stably transfected with hACE2 were seeded in 96-well culture plates at a density of 5,000 cells per well. Immune sera were then diluted with pseudo-virus containing supernatant and incubated at 37 °C for 1 h and then transferred to the target cells. After overnight incubation, fresh cell culture medium was changed and the cells were further cultured for two days. Luciferase activity was then analyzed using luciferase assay substrate (Promega, Cat# E1483).

### NHP PBMCs stimulation, intracellular staining and flow cytometry

The monkey whole blood samples were collected before (week 0) or after (week 8) AAV-ie-S1 vaccination, PBMCs were isolated using Ficoll (GE, Cat# 17144002) gradient centrifugation following the manufacturer’s instruction. PBMCs were cultured in RPMI 1640 medium (Gibco, Cat# 11875119) supplemented with 10% heat inactivated fetal bovine serum (HIFBS) (Gibco, Cat# 10091148), and penicillin/streptomycin. Cells were stimulated with synthetic SARS-CoV-2 S peptides pool (Genscript, Cat# RP30020) at the concentration of 2 μg/mL for 12 h and then incubated with 5 μg/mL Brefeldin A (MCE, Cat# HY-16592). Cells stimulated with 50 ng/mL PMA (Sigma, Cat# P8139) and 1 μM ionomycin (J&K, Cat# 464833) for 4 h were used as positive control.

After stimulation, cells were washed with PBS and stained using LIVE/DEAD fixable aqua (Thermo Fisher Scientific, Cat# L34957) to exclude dead cells. Subsequently, cells were stained using pacific blue conjugated anti-human CD3 (BD Bioscience, Cat# 558124), FITC conjugated anti-human CD4 (BD Bioscience, Cat# 550628), PE conjugated anti-human CD8 (BD Bioscience, Cat# 557086). Intracellular staining of the cytokines was performed after fixation and permeabilization using APC conjugated anti-human IFNγ (BD Bioscience, Cat# 551385), anti-human IL2 (BD Bioscience, Cat# 551383), and anti-human TNFα (BD Bioscience, Cat# 551384) accordingly. The percentages of cytokine positive CD4^+^ or CD8^+^ cells were analyzed using flow cytometer (Cytoflex S, Beckman).

### S-binding assay using FACS

The plasmids encoding wild type SARS-CoV-2 or circulating variants (B.1.351 and B.1.1.7) S proteins with a C-terminal fused GFP were transfected into HEK293T cells. After 48 h culture, cells were dissociated and incubated with serially diluted monkey immune sera at 4 °C for 1h. After PBS washing, cells were further stained using alexa-555 conjugated donkey anti-human IgG (Thermo Fisher Scientific, Cat# A21433). After extensive washing, the percentages of anti-human IgG postive cells in GFP positive cells were then analyzed using flow cytometer (Cytoflex S, Beckman). LIVE/DEAD fixable aqua (Thermo Fisher Scientific, Cat# L34957) was used to exclude dead cells.

### Statistics

Antibody titers of ELISA assay, EC_50_ value of pseudo-virus neutralization assay were determined using non-linear regression analysis using Graphpad PRISM. Data are shown as mean ± SD. Numbers of replicates for experiments are described in the figure legends.

## Acknowledgements

We thank the staff from the Bio-imaging core and Mammalian core of iHuman Institute, ShanghaiTech University and the High Throughput Screening platform of Shanghai Institute for Advanced Immunochemical Studies (SIAIS), ShanghaiTech University for their technical assistance. This work was supported by ShanghaiTech, and grants from the National Key Research and Development Program of China (2017YFC1001301 (G.Z.)) and National Natural Science Foundation of China (31871487(C.L.)).

## Author contributions

G.Z., S.Z., and F.Z. conceived and designed the experiments. S.Z., F.Z., and J.K. designed the vaccine candidates, and performed ELISA, pseudovirus neutralization assays. J.Y. performed protein purification and immunofluorescence experiment. All the authors contributed to data analysis, interpretation, and presentation. G.Z. and S.Z. wrote the manuscript with contributions from all the authors.

---

**Extended Data Fig. 1.**
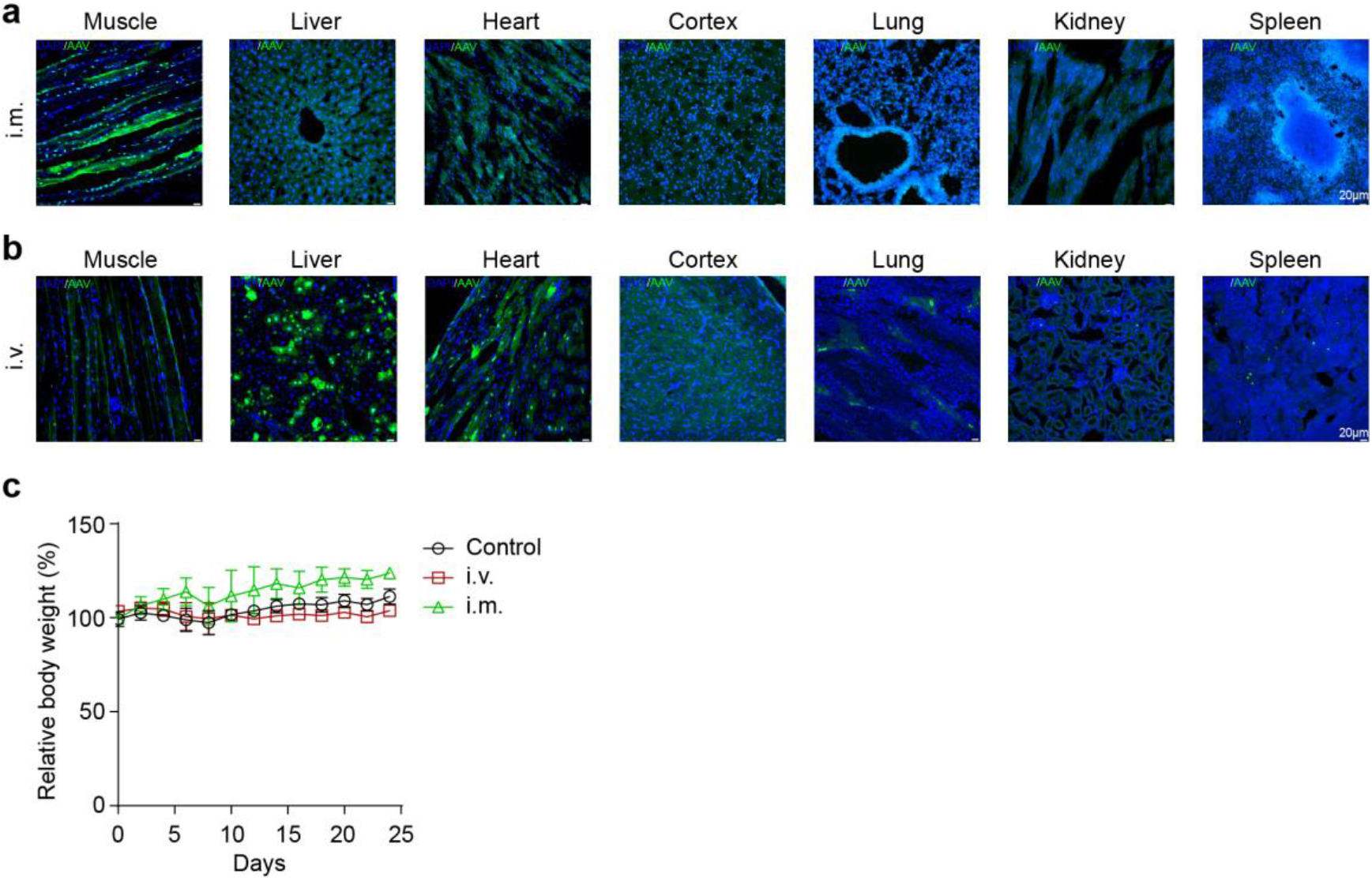
Tissue tropism of AAV-ie. Representative fluorescence imaging of indicated organ slices after i.m. (**a**) and i.v. (**b**) injection of AAV-ie-GFP. Scale bar, 20 μm. **c**, Relative body weight of mice after i.m. or i.v. injection of AAV-ie-GFP. N = 3 in each group.

**Extended Data Fig. 2.**
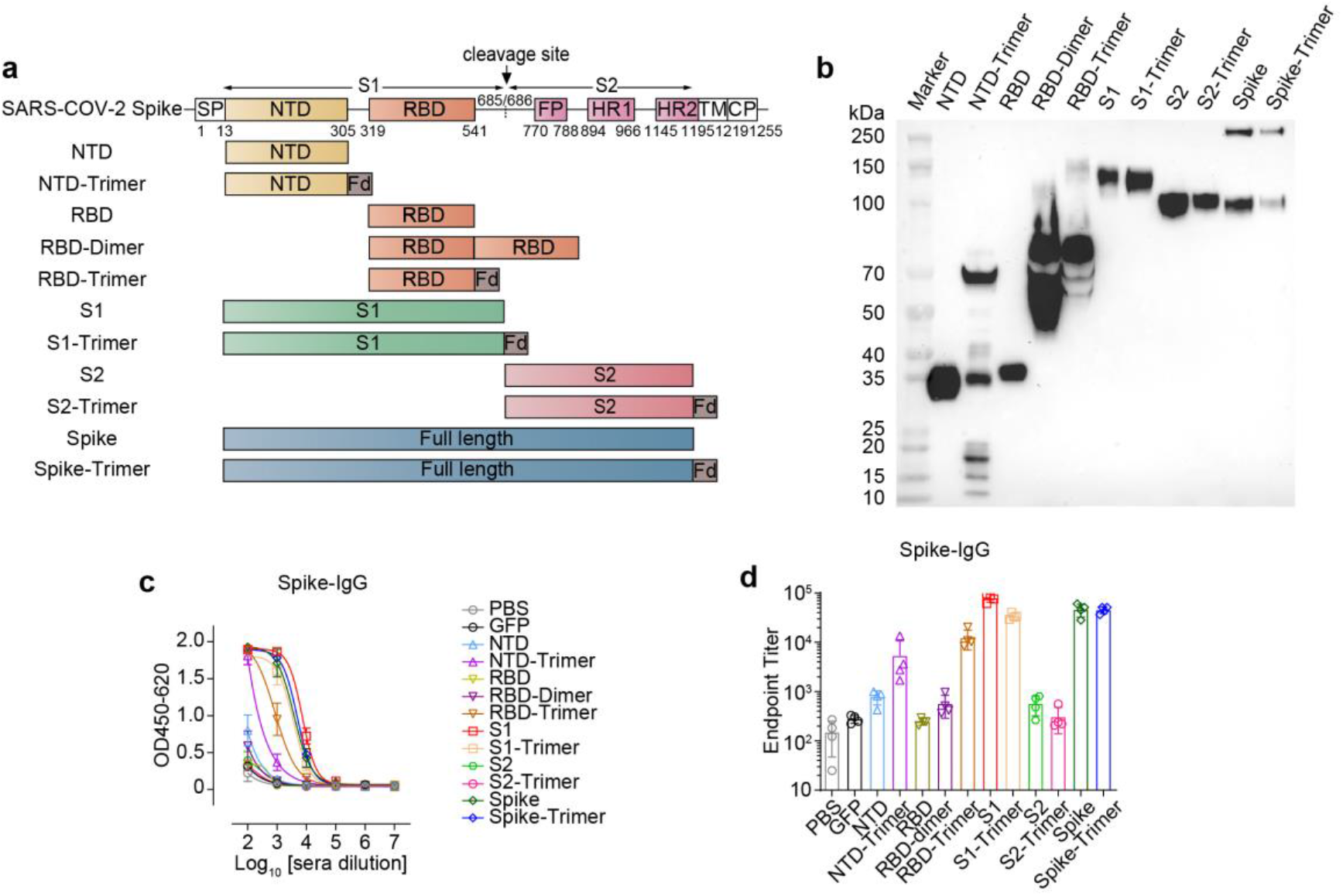
Systematic SARS-CoV-2 S antigen screening. **a**, AAV-ie vectored vaccines design. **b**, Detection of S antigens expression using Western blot in HEK293T cells infected by related plasmids. **c**, S-Binding IgG antibody titers of mice sera after vaccination (i.v.) with indicated antigens at week 2, PBS and AAV-ie-GFP were used as negative control. Experiments were performed in triplicates. **d**, End-point titers summary in **c**, N = 4 in each group,

**Extended Data Fig. 3.**
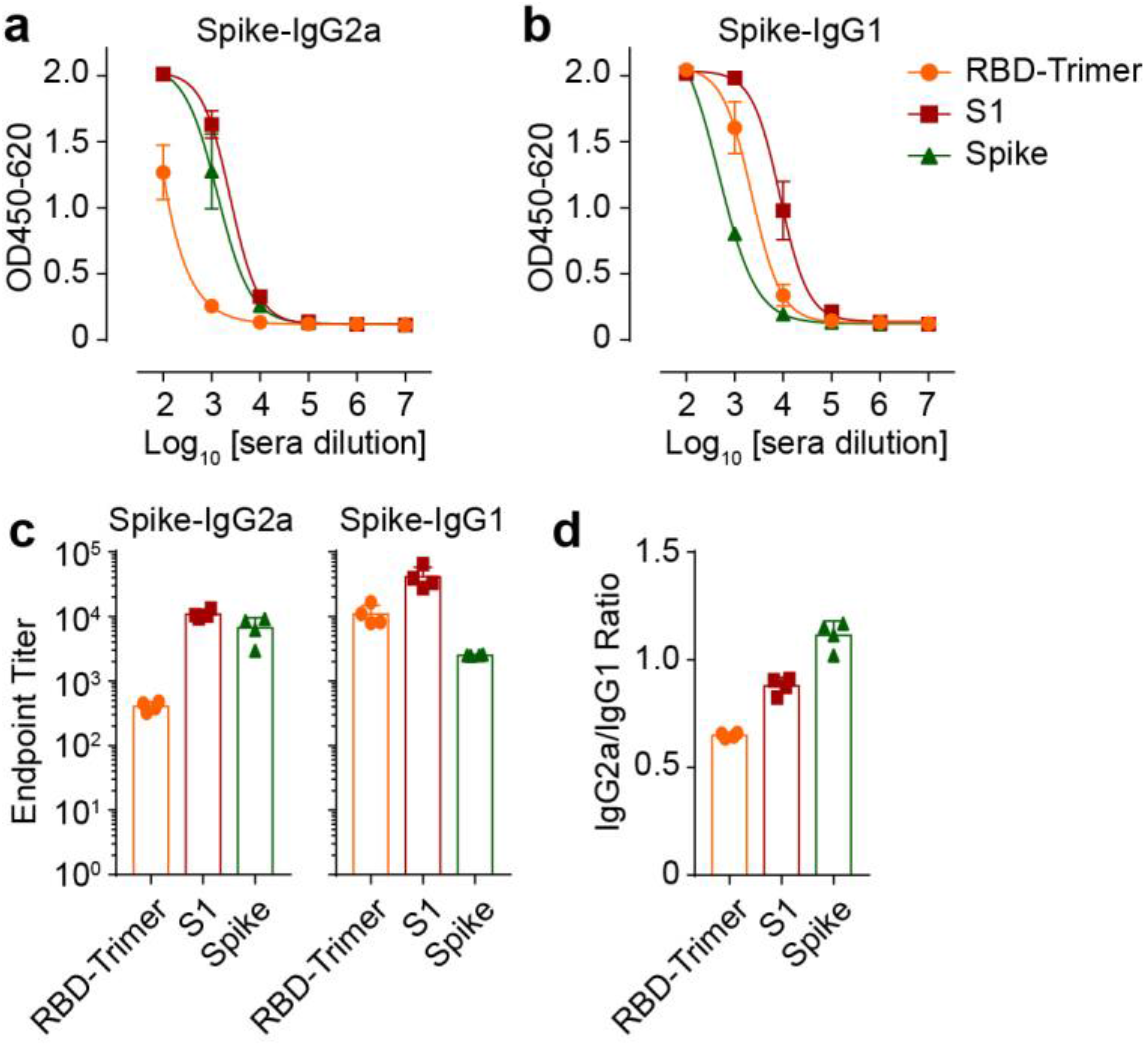
SARS-CoV-2 S-Binding antibody isotype titers of vaccinated sera in mice. **a-b**, Determination of IgG2a (**a**) and IgG1 (**b**) titers using ELISA assay. Experiments were performed in triplicates. **c**, Summary of the endpoint titers of IgG2a and IgG1 in indicated immune sera (N = 4, each with 3 replicates). **d**, End-point titer ratios of IgG2a to IgG1 of three vaccines.

**Extended Data Fig. 4.**
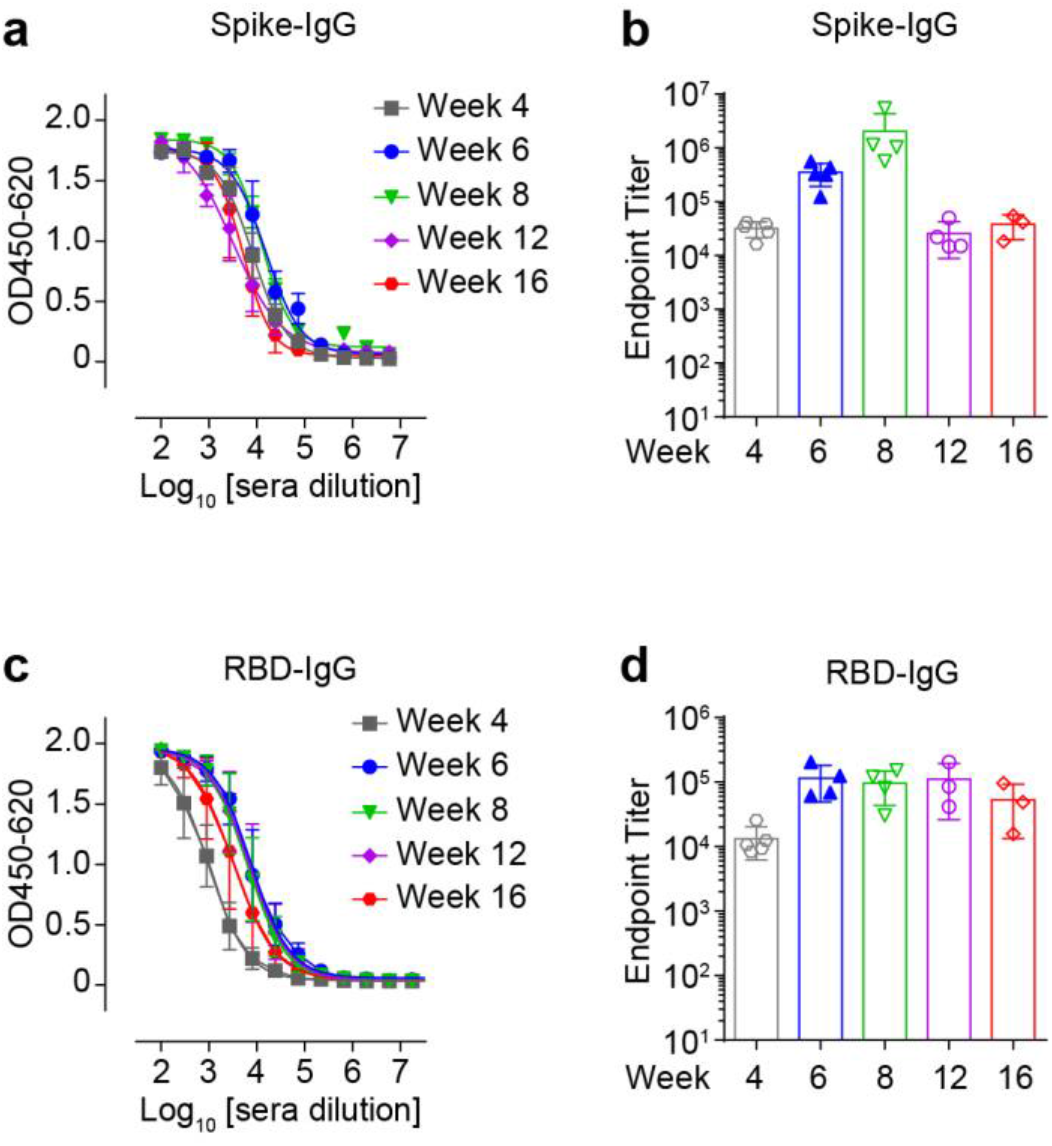
Monitoring of SARS-CoV-2 S- and RBD-Binding antibody titers of immune sera in AAV-ie-S1 vaccinated mice. **a-b**, S-Binding IgG antibody titers of mice sera after vaccination (i.m.) with AAV-ie-S1 at indicated time. **a**, Determination of S-binding IgG antibody titers using ELISA assay. **b**, Endpoint titers of S-binding IgG antibody at indicated time (N = 3 - 5, each with 3 replicates). **c-d**, S-Binding IgG antibody titers of mice sera after vaccination (i.m.) with AAV-ie-S1 at indicated time. **c**, Determination of RBD-binding IgG antibody titers. **d**, Endpoint titers of RBD-binding IgG antibody (N = 3 - 5, each with 3 replicates).

**Extended Data Fig. 5.**
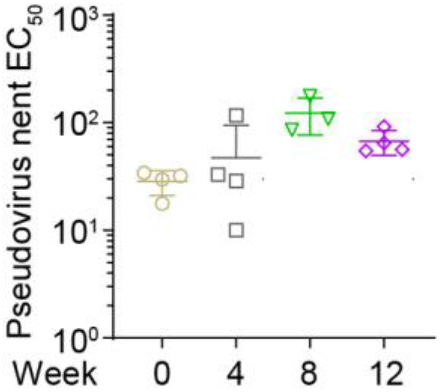
SARS-CoV-2 pseudo-virus neutralizing EC_50_ values of AAV-ie-S1 immune sera at indicated times. (N = 3 - 5, each with 3 replicates).

**Extended Data Fig. 6.**
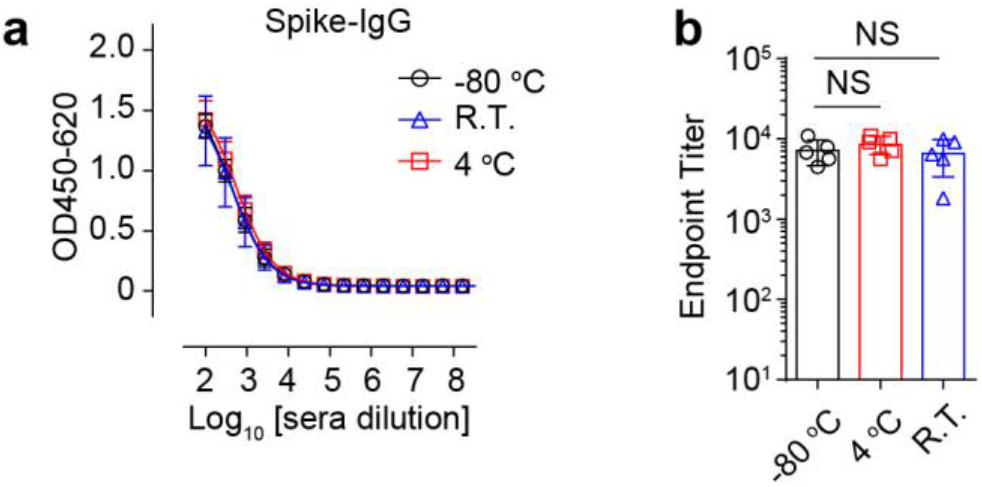
Thermostability of AAV-ie-S1. **a**, S-Binding IgG antibody titer determination of mice sera using ELISA on week 2 after vaccination (i.m.) with AAV-ie-S1 kept at indicated temperatures for 2 weeks. Experiments were performed in triplicates. **b**, Summary of Endpoint titers. NS: no significance. (N = 5 in each group)

**Extended Data Fig. 7.**
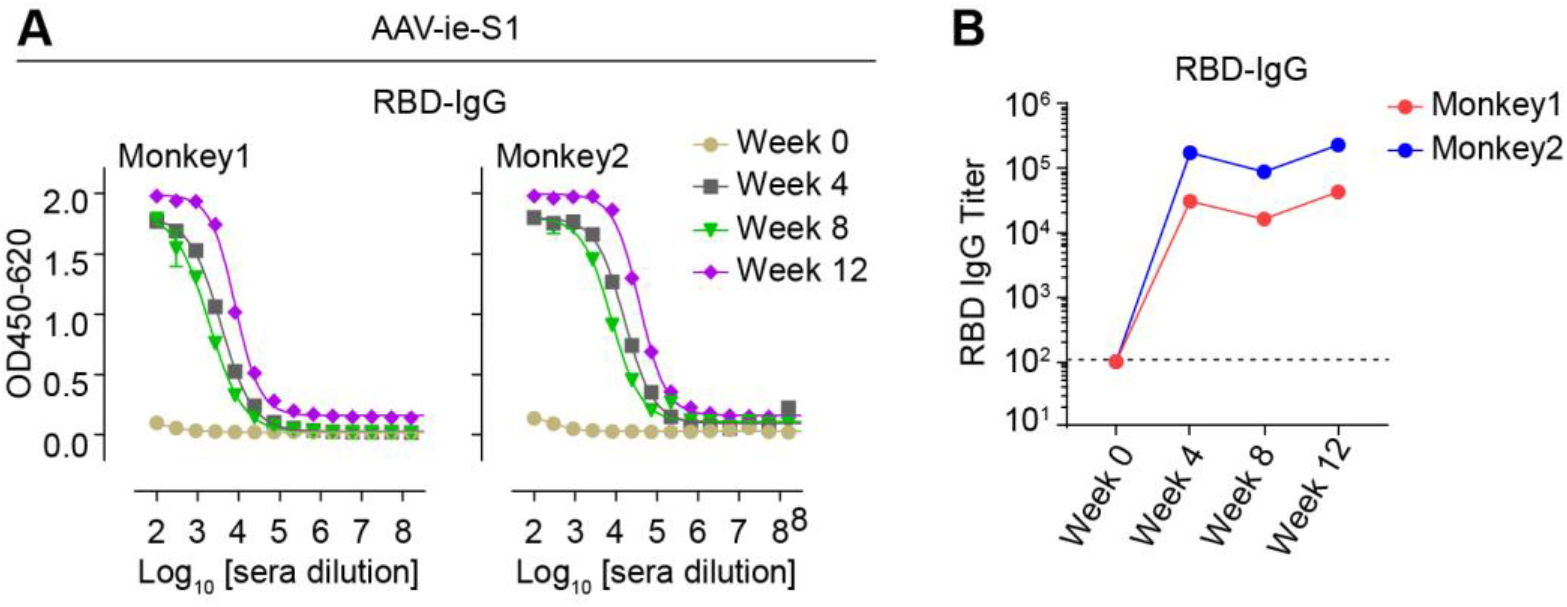
Monitoring of SARS-CoV-2 RBD-Binding antibody titers of immune sera in AAV-ie-S1 vaccinated NHPs. **A**, Determination of RBD-binding IgG antibody titers using ELISA assay. Experiments were performed in triplicates. **B**, Endpoint titers of RBD-binding IgG antibody at indicated time.

**Extended Data Fig. 8.**
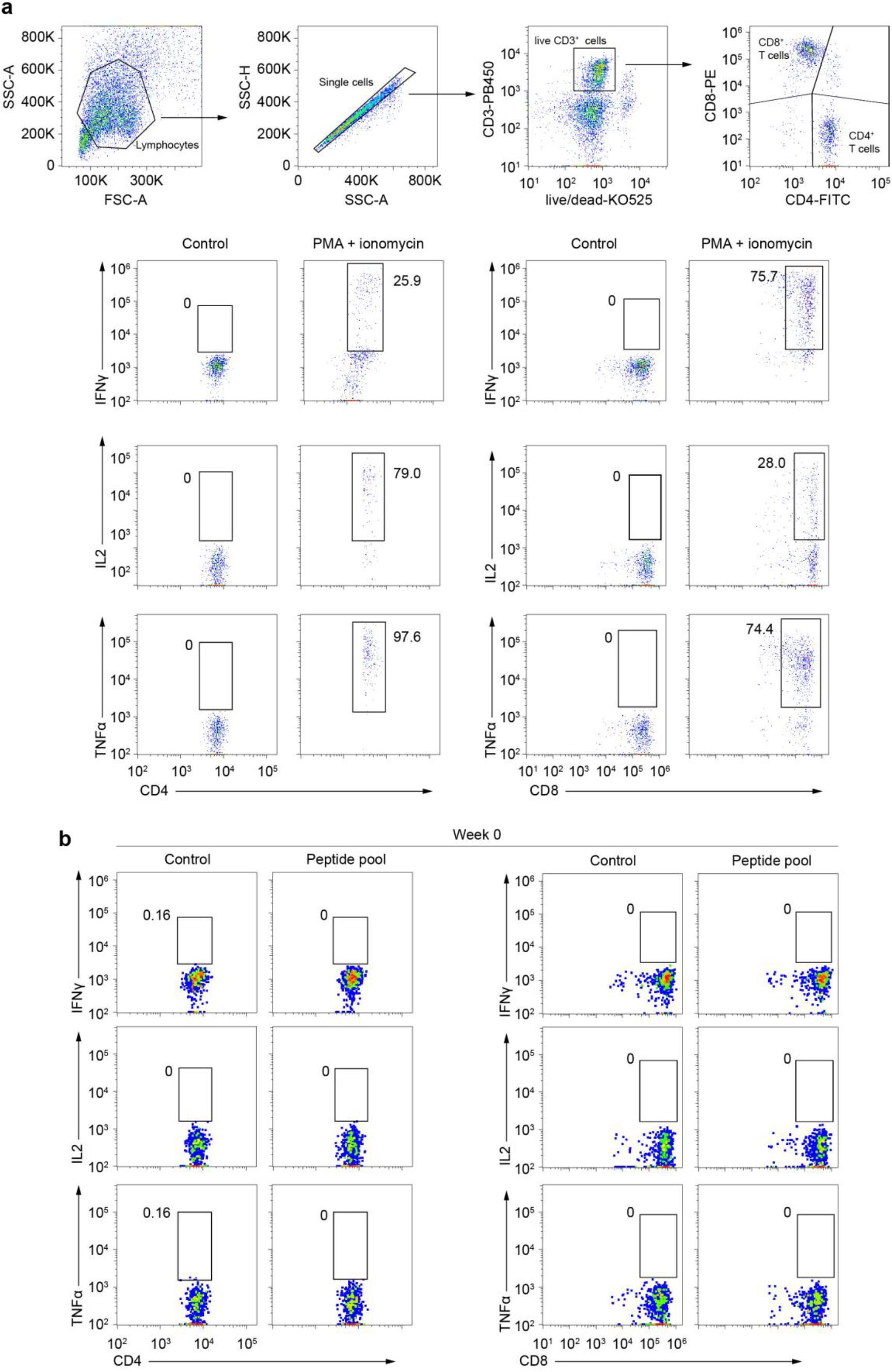
Flow cytometry analysis of SARS-CoV-2 S-specific T cells in NHPs. **a**, Gating strategy. PBMCs stimulated with PMA and ionomycin were used as positive controls. **b**, Gating summary of SARS-CoV-2 S-specific CD4^+^ and CD8^+^ T cells before vaccination (Week 0).

**Extended Data Fig. 9.**
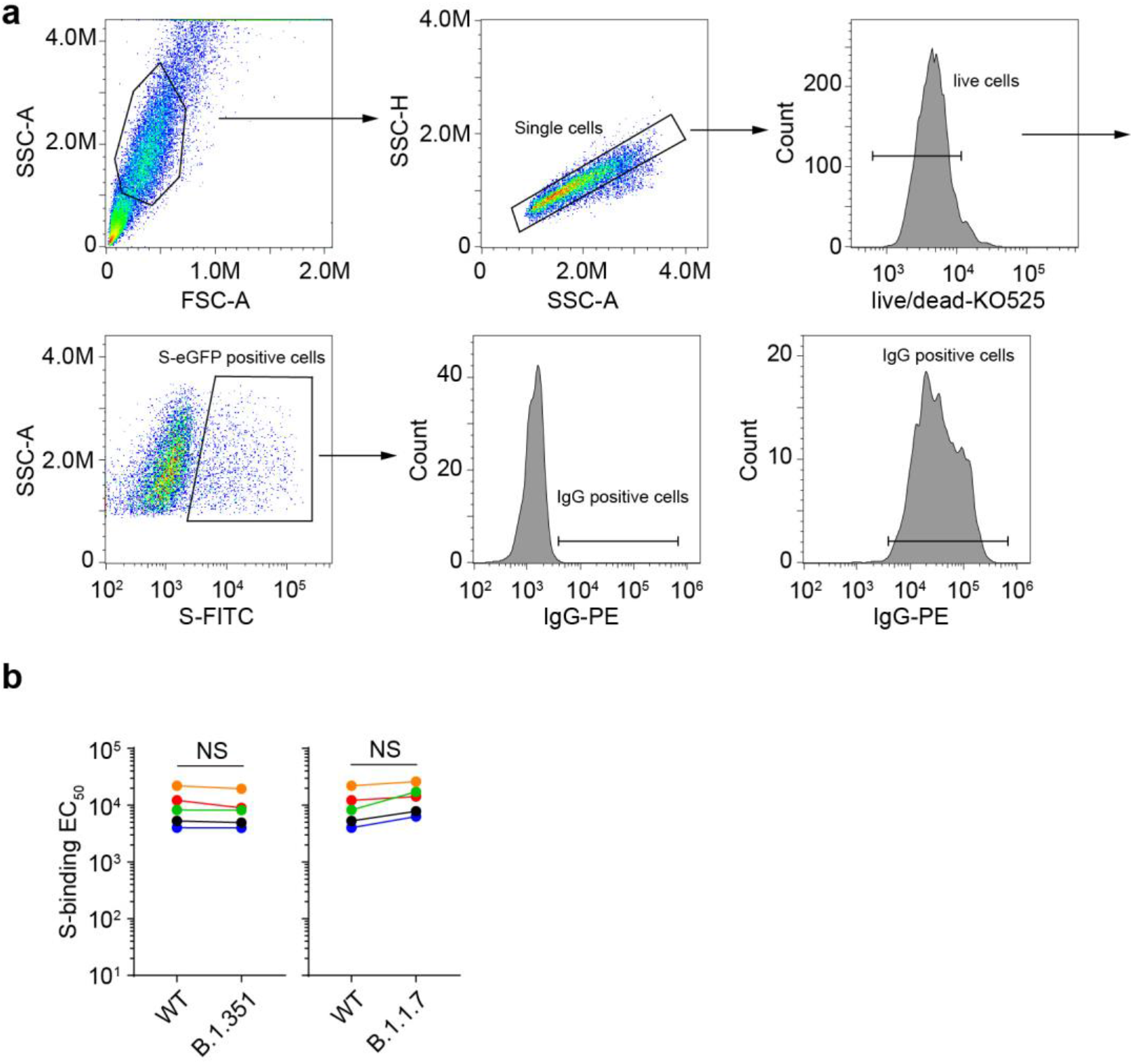
FACS-based S-binding Assay. **a,** Gating strategy of FACS based SARS-CoV-2 S-Binding assay. **b,** S-binding EC_50_ values change of mouse immune sera against B.1.351 and B.1.1.7. N = 5 in each group.

## References

1 Xia, S. et al. Safety and immunogenicity of an inactivated SARS-CoV-2 vaccine, BBIBP-CorV: a randomised, double-blind, placebo-controlled, phase 1/2 trial. The Lancet Infectious Diseases 21, 39–51, doi:10.1016/s1473-3099(20)30831-8 (2021).

2 Wu, Z. et al. Safety, tolerability, and immunogenicity of an inactivated SARS-CoV-2 vaccine (CoronaVac) in healthy adults aged 60 years and older: a randomised, double-blind, placebo-controlled, phase 1/2 clinical trial. The Lancet Infectious Diseases, doi:10.1016/s1473-3099(20)30987-7 (2021).

3 Zhang, Y. et al. Safety, tolerability, and immunogenicity of an inactivated SARS-CoV-2 vaccine in healthy adults aged 18–59 years: a randomised, double-blind, placebo-controlled, phase 1/2 clinical trial. The Lancet Infectious Diseases, doi:10.1016/s1473-3099(20)30843-4 (2020).

4 Polack, F. P. et al. Safety and Efficacy of the BNT162b2 mRNA Covid-19 Vaccine. N Engl J Med, doi:10.1056/NEJMoa2034577 (2020).

5 Baden, L. R. et al. Efficacy and Safety of the mRNA-1273 SARS-CoV-2 Vaccine. N Engl J Med 384, 403–416, doi:10.1056/NEJMoa2035389 (2021).

6 Yang, J. et al. A vaccine targeting the RBD of the S protein of SARS-CoV-2 induces protective immunity. Nature, doi:10.1038/s41586-020-2599-8 (2020).

7 Ramasamy, M. N. et al. Safety and immunogenicity of ChAdOx1 nCoV-19 vaccine administered in a prime-boost regimen in young and old adults (COV002): a single-blind, randomised, controlled, phase 2/3 trial. The Lancet 396, 1979–1993, doi:10.1016/s0140-6736(20)32466-1 (2020).

8 Zhu, F.-C. et al. Immunogenicity and safety of a recombinant adenovirus type-5-vectored COVID-19 vaccine in healthy adults aged 18 years or older: a randomised, double-blind, placebo-controlled, phase 2 trial. The Lancet 396, 479–488, doi:10.1016/s0140-6736(20)31605-6 (2020).

9 Chen, R. E. et al. Resistance of SARS-CoV-2 variants to neutralization by monoclonal and serum-derived polyclonal antibodies. Nat Med, doi:10.1038/s41591-021-01294-w (2021).

10 Wang, Z. et al. mRNA vaccine-elicited antibodies to SARS-CoV-2 and circulating variants. Nature, doi:10.1038/s41586-021-03324-6 (2021).

11 Wang, P. et al. Antibody Resistance of SARS-CoV-2 Variants B.1.351 and B.1.1.7. Nature, doi:10.1038/s41586-021-03398-2 (2021).

12 Garcia-Beltran, W. F. et al. Multiple SARS-CoV-2 variants escape neutralization by vaccine-induced humoral immunity. Cell, doi:10.1016/j.cell.2021.03.013 (2021).

13 Wang, G.-L. et al. Susceptibility of Circulating SARS-CoV-2 Variants to Neutralization. N Engl J Med, doi:10.1056/NEJMc2103022 (2021).

14 Li, C. & Samulski, R. J. Engineering adeno-associated virus vectors for gene therapy. Nat Rev Genet 21, 255–272, doi:10.1038/s41576-019-0205-4 (2020).

15 Demminger, D. E. et al. Adeno-associated virus-vectored influenza vaccine elicits neutralizing and Fcgamma receptor-activating antibodies. EMBO Mol Med 12, e10938, doi:10.15252/emmm.201910938 (2020).

16 Vardas, E. et al. A phase 2 study to evaluate the safety and immunogenicity of a recombinant HIV type 1 vaccine based on adeno-associated virus. AIDS Res Hum Retroviruses 26, 933–942, doi:10.1089/aid.2009.0242 (2010).

17 Tan, F. et al. AAV-ie enables safe and efficient gene transfer to inner ear cells. Nat Commun 10, 3733, doi:10.1038/s41467-019-11687-8 (2019).

18 Kroger, A., Bahta, L. & Hunter, P. General Best Practice Guidelines for Immunization: Best Practices Guidance of the Advisory Committee on Immunization Practices (ACIP). (2020).

19 Zabaleta, N. et al. Immunogenicity of an AAV-based, room-temperature stable, single dose COVID-19 vaccine in mice and non-human primates. bioRxiv, doi:10.1101/2021.01.05.422952 (2021).

20 Wrapp, D. et al. Cryo-EM structure of the 2019-nCoV spike in the prefusion conformation. Science 367, 1260, doi:10.1126/science.abb2507 (2020).

